# Small molecule telomerase inhibitors are also potent inhibitors of telomeric C-strand synthesis

**DOI:** 10.1101/2024.04.01.587626

**Authors:** Kaitlin Johnson, Julia M. Seidel, Thomas R. Cech

## Abstract

Telomere replication is essential for continued proliferation of human cells, such as stem cells and cancer cells. Telomerase lengthens the telomeric G-strand, while C-strand replication is accomplished by CST-polymerase α -primase (CST-PP). Replication of both strands is inhibited by formation of G-quadruplex (GQ) structures in the G-rich single-stranded DNA. TMPyP4 and pyridostatin (PDS), which stabilize GQ structures in both DNA and RNA, inhibit telomerase *in vitro*, and they cause telomere shortening in human cells that has been attributed to telomerase inhibition. Here, we show that TMPyP4 and PDS also inhibit C-strand synthesis by stabilizing DNA secondary structures and thereby preventing CST-PP from binding to telomeric DNA. We also show that these small molecules inhibit CST-PP binding to a DNA sequence containing no consecutive guanine residues, which is unlikely to form GQs. Thus, while these “telomerase inhibitors” indeed inhibit telomerase, they are also robust inhibitors of telomeric C-strand synthesis. Furthermore, given their limited specificity for GQ structures, they may disrupt many other protein-nucleic acid interactions in human cells.

## INTRODUCTION

Telomeric DNA is a guanine-rich repeating sequence ((TTAGGG)_n_ in humans) at the natural ends of linear chromosomes, with the double-stranded DNA repeats terminating in a 3’ single-stranded overhang (Lim and Cech 2021). These ends cannot be replicated by standard polymerases, leading to the progressive shortening of telomeres over time (Harley et al. 1990). Some highly proliferating cells (e.g., human stem cells, germline cells and most cancer cells) overcome this problem by producing telomerase, a ribonucleoprotein complex with the unique ability to extend the 3’ end of the telomeric G-strand (Kim et al. 1994). The CST (CTC1-STN1-TEN1) protein complex regulates telomere synthesis, both preventing re-initiation of G-strand synthesis by telomerase and directing C-strand synthesis through its direct association with telomeric ssDNA and polymerase α-primase (Martin et al. 2000; Qi and Zakian 2000; Miyake et al. 2009; Chen et al. 2012; Wang et al. 2012; Lue et al. 2014). Recent cryo-EM structures and biochemical reconstitution have continued to illuminate details of telomeric C-strand synthesis (Cai et al. 2022; He et al. 2022; Zaug et al. 2022).

Secondary structures within single-stranded DNA pose a challenge to telomere replication. G-rich DNA sequences, including the repeating human telomeric sequence (TTAGGG)_n_, are capable of forming G-quadruplexes, structures in which four guanines interact to form a tetrad or “G-quartet”, and several such quartets stack to form a quadruplex (Williamson et al. 1989; Spiegel et al. 2020). These stable structures are inhibitory to many enzymatic processes, including telomerase activity; GQ formation can inhibit telomerase both by preventing its initial association with the telomere tail and by forming within the telomere-protein complex during extension (Zahler et al. 1991; Jansson et al. 2019). GQs can also form in the open promoters of several oncogenes (Simonsson et al. 1998; Cogoi and Xodo 2006). Thus, specific stabilization of GQs by small molecules could prove to be a powerful tool for cancer therapeutics (Kosiol et al. 2021).

Some telomere-associated proteins, including POT1 (Protection of Telomeres 1) (Zaug et al. 2005), can disrupt GQs on the telomere tail to restore telomerase access. CST has also been shown to resolve GQs *in vitro* (Bhattacharjee et al. 2017; Zhang et al. 2019) and through cooperation with the helicase RECQ4 *in vivo* (Li et al. 2023). It is recruited to stalled replication forks throughout the genome to resolve GQs and restart replication (Stewart et al. 2012). CST-PP can also participate in double strand break repair, filling in resected 3’ ends to enable homology-directed repair (Mirman et al. 2018).

G-quadruplex stabilizers including TMPyP4 and PDS are often called “telomerase inhibitors,” and certainly they inhibit telomerase action *in vitro* (Wheelhouse et al. 1998; De Cian et al. 2007; Rodriguez et al. 2008). In cells, these and other small molecule GQ-stabilizers lead to progressive telomere shortening and apoptosis (Fujimori et al. 2011; Müller et al. 2010), but telomere dysfunction may not be limited to inhibition of telomerase; other factors may include increased DNA damage at telomeres (Rodriguez et al. 2008; Fujimori et al. 2011) and downregulation of telomeric protein expression (Gomez et al. 2010). Furthermore, it is unknown whether the demonstrated impact of these small molecules on telomere extension is exclusive to G-strand synthesis by telomerase, or if they also inhibit C-strand replication by CST-PP. We previously demonstrated that CST-PP activity is enhanced ten-fold when GQs are not present on the template (Zaug et al. 2022), indicating that although CST-PP effectively resolves such structures, they do indeed pose a barrier. It is unknown whether CST-PP would be able to overcome further stabilization of GQs by small molecules.

In this study, we report the inhibition of human CST-PP activity by TMPyP4 and PDS. We found that C-strand synthesis was inhibited by both small molecules in the presence of GQ-stabilizing cations Na^+^ and K^+^, but – unexpectedly – also in the presence of Li^+^ or with template DNA that contained no consecutive G nucleotides. We demonstrated that the inhibition of synthesis is caused by CST-PP’s inability to bind to the templates due to secondary structures, in some cases unpredicted, that are highly stabilized by the small molecules. These results highlight the need for caution and controls when utilizing GQ-stabilizing drugs, particularly in the complex environment of cells. This work emphasizes the importance of considering GQ ligands not as telomerase inhibitors, but as DNA binders that more generally disrupt enzyme activity.

## RESULTS

We first confirmed telomerase inhibition by TMPyP4 and PDS under our standard assay conditions in vitro by incubating telomerase, dNTPs, and 50 nM 3xTEL DNA primer (1xTEL = TTAGGG) with increasing concentrations of each small molecule at 30 °C **(Supplemental Fig. S1A**). TMPyP4 and PDS gave half maximal inhibitory concentrations (IC50) of 6400 and 520 nM, respectively (**Supplemental Fig S1B)**; these values are in general agreement with previously published IC50s under other conditions (Wheelhouse et al. 1998; De Cian et al. 2007; Rodriguez et al. 2008).

### TMPyP4 inhibits CST-PP synthesis

Telomeric C-strand synthesis was performed with CST-PP purified from cultured human cells (HEK293T), a synthetic telomeric DNA template (9xTEL, (TTAGGG)_9_), ribonucleotides to allow primase activity, and deoxynucleotides including α -^32^P-dATP to monitor polymerase α extension. To test whether TMPyP4 would inhibit DNA synthesis by CST-PP, we preincubated 25 nM 9xTEL with increasing concentrations of the drug, then added 25 nM CST-PP and nucleotides and observed C-strand synthesis by gel electrophoresis **(Fig. 1A, left**). To serve as a size marker and recovery control, a small fraction of the template (T) was radiolabeled. The expected synthesis products were observed, all smaller than the 54 nt template **(Supplemental Fig S2A)**. The limited extension reflects the limited processivity of CST-PP, and the six-nucleotide ladder of products is due to multiple initiation sites on the six-nucleotide repeat template (Zaug et al. 2022).

**FIGURE 1.**
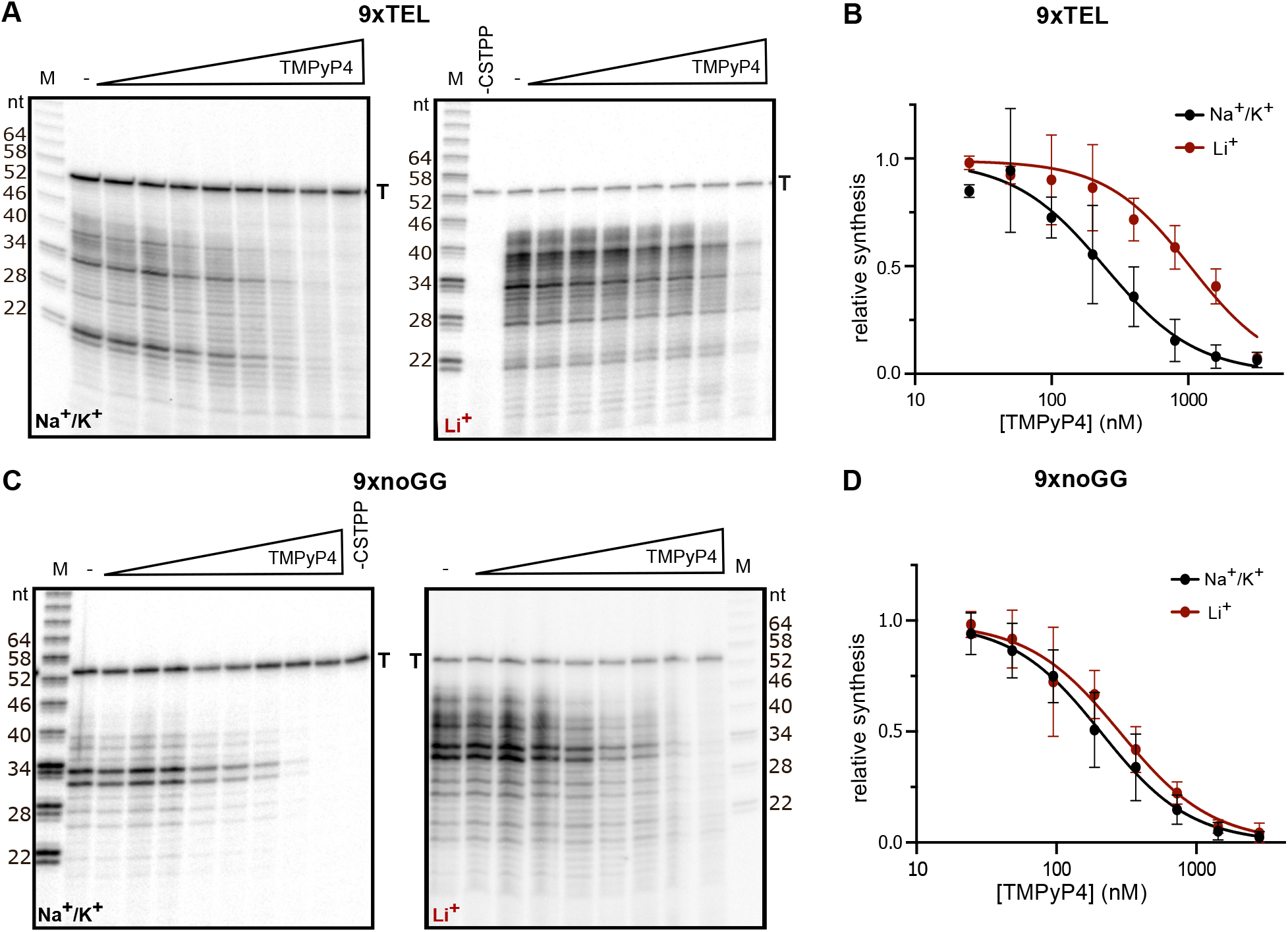
Inhibition of CST-polymerase α-primase activity by TMPyP4. (A) Representative gels showing C-strand synthesis on the 9xTEL template in NaCl/KCl (left) or LiCl (right) with trace amount of radiolabeled template (T) serving as recovery control. Two-fold dilutions of TMPyP4 (highest concentration: 3200 nM) were incubated with template prior to addition of CST-PP and nucleotides. (B) Plot of relative synthesis intensities as a function of TMPyP4 concentration in NaCl/KCl or LiCl. Points represent mean values and error bars represent SD for n = 5 (Na^+^/K^+^) or n = 4 (Li^+^) independent experiments. (C,D) Same as panels A, B except that template was 9xnoGG. For panel D, n = 4.

Normalizing the intensity of products at each concentration of TMPyP4 to the no-drug control gave a half maximal inhibitory concentration of TMPyP4 for synthesis (IC50_synthesis_) of 250 ± 63 nM **(Fig. 1B**, **Table 1).** To determine if inhibition was dependent on GQ structures in the template DNA, we repeated the experiment, replacing the standard NaCl and KCl in the reaction with LiCl to destabilize quadruplex formation **(Fig. 1A, right)**. We found that TMPyP4 still inhibited synthesis, but the IC50_synthesis_ was about 4-fold higher in Li^+^ than in Na^+^/K^+^ **(Fig. 1B**, **Table 1).** Thus, TMPyP4 inhibits CST-PP activity on a GQ-forming template, and inhibition is more efficient in GQ-stabilizing conditions.

**TABLE 1.**
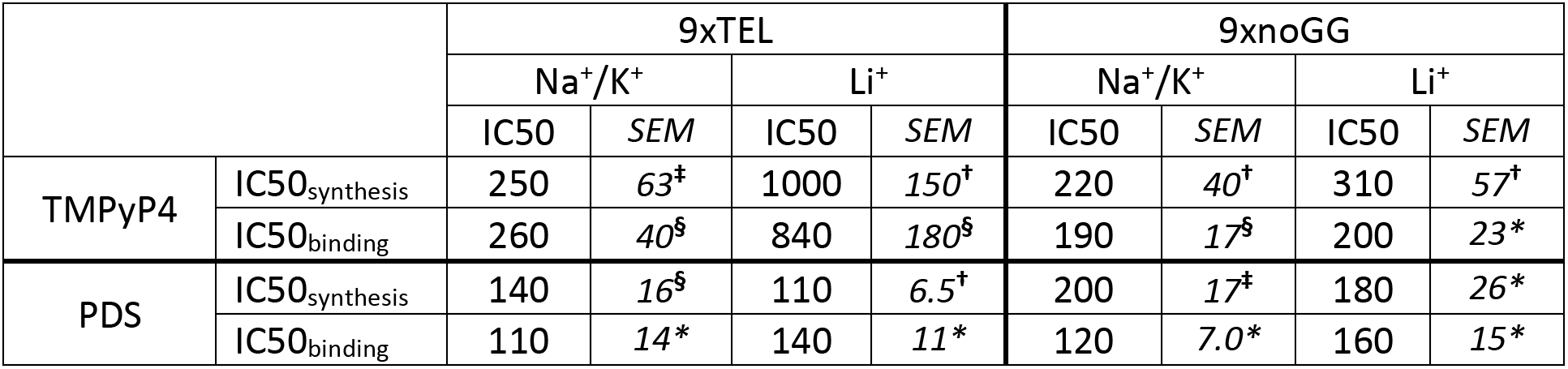
IC50 values (nM) of TMPyP4 or PDS inhibition of CST-PP synthesis or binding to the DNA template indicated. IC50 and SEM calculated from n = 3 (*), 4 (†), 5 (‡), or 6 (§) independent replicates as indicated.

|  |  | 9xTEL |  |  |  | 9xnoGG |  |  |  |
| --- | --- | --- | --- | --- | --- | --- | --- | --- | --- |
|  |  | Na<sup>+</sup>/K<sup>+</sup> |  | Li<sup>+</sup> |  | Na<sup>+</sup>/K<sup>+</sup> |  | Li<sup>+</sup> |  |
|  |  | IC50 | SEM | IC50 | SEM | IC50 | SEM | IC50 | SEM |
| TMPyP4 | IC50<sub>synthesis</sub> | 250 | 63<sup>‡</sup> | 1000 | 150<sup>†</sup> | 220 | 40<sup>†</sup> | 310 | 57<sup>†</sup> |
|  | IC50<sub>binding</sub> | 260 | 40<sup>§</sup> | 840 | 180<sup>§</sup> | 190 | 17<sup>§</sup> | 200 | 23<sup>*</sup> |
| PDS | IC50<sub>synthesis</sub> | 140 | 16<sup>§</sup> | 110 | 6.5<sup>†</sup> | 200 | 17<sup>‡</sup> | 180 | 26<sup>*</sup> |
|  | IC50<sub>binding</sub> | 110 | 14<sup>*</sup> | 140 | 11<sup>*</sup> | 120 | 7.0<sup>*</sup> | 160 | 15<sup>*</sup> |

In Na^+^/K^+^, the most prominent product with or without drug was around 21 nt **(Fig. 1A, left).** A 21 nt product could be the result of GQ formation at either end of the template that essentially limits the available length of the template and therefore of the product; a GQ at the 3’ end could bias initiation at a central repeat rather than the end, or a GQ at the 5’ end may hinder extension by CST-PP **(Supplemental Fig. S2B)**. Notably, the distribution of product sizes increased in Li^+^ compared to the GQ-stabilizing condition and the ∼21 nt product was much less intense; addition of TMPyP4 did not lead to accumulation of smaller products **(Fig. 1A, right)**.

To further examine whether inhibition by TMPyP4 was GQ-dependent, we tested a mutated sequence that contained no consecutive G’s [(TGAGTG)_9_ which we call 9xnoGG]. We found that TMPyP4 unexpectedly inhibited synthesis from this template **(Fig. 1C)** with about the same IC50_synthesis_ as that of 9xTEL under the same conditions **(Fig. 1D**, **Table 1)**. Unlike 9xTEL, the IC50_synthesis_ in Li^+^ was similar to the IC50 in Na^+^/K^+^. Thus, TMPyP4 inhibits CST-PP activity on templates without consecutive G’s in a cation-independent manner.

### Inhibition by TMPyP4 occurs at the level of CST-PP binding to template DNA

TMPyP4 inhibited all extension products to a similar extent, suggesting that inhibition was more at the level of initiation than elongation. To test directly if TMPyP4 inhibited CST-PP by preventing association with the templates, we next measured the effect of the drug on the proportion of DNA bound by CST-PP. We used the same conditions as in the synthesis reaction and observed that the DNA was about half bound in the absence of inhibitor. We prepared DNA templates with an increasing concentration of TMPyP4 as before, then incubated with CST-PP and analyzed by EMSA (Electrophoretic Mobility Shift Assay) **(Fig. 2A)**. By normalizing the fraction of DNA bound at each concentration of inhibitor to fraction bound without drug, we determined IC50 values for binding (IC50_binding_) in the two salts for both 9xTEL and 9xnoGG **(Fig. 2B**, **Table 1)**. The four IC50_binding_ values are in agreement with the respective IC50_synthesis_, indicating that TMPyP4 inhibits synthesis by preventing CST-PP from associating with DNA templates.

**FIGURE 2.**
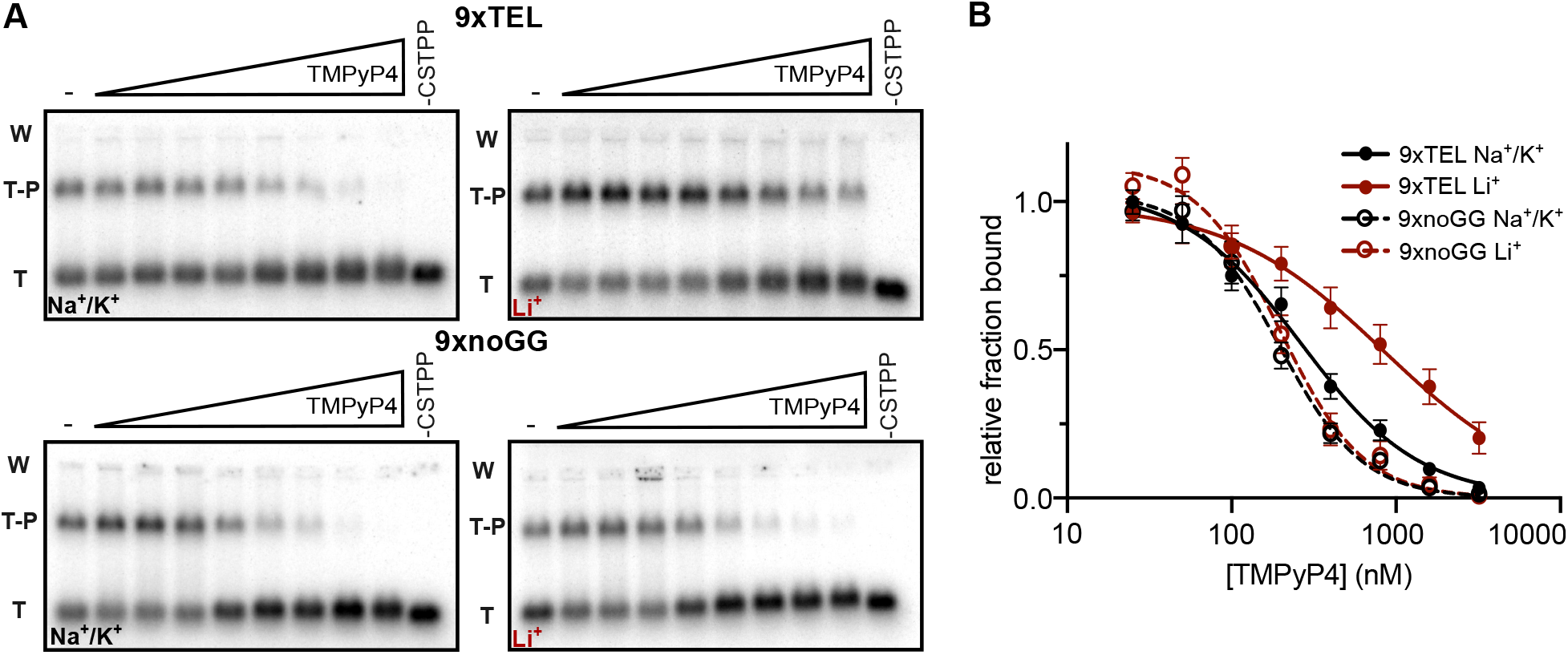
TMPyP4 inhibits CST-polymerase α-primase binding to template DNA. (A) Representative gels of CST-PP binding to radiolabeled template DNA in NaCl/KCl (left) or LiCl (right) following incubation of DNA with increasing concentrations of TMPyP4 (two-fold dilutions from 3200 nM). T = Template, T-P = Template-Protein complex, W = Well. (B) Plot of fraction DNA bound at each concentration of TMPyP4 normalized to fraction bound with no inhibitor in NaCl/KCl or LiCl. Points represent mean values and error bars represent SD for n = 6 independent experiments, except 9xnoGG in Li^+^ (n = 4).

### Pyridostatin inhibits CST-Pol α-primase activity by inhibiting binding to the template

Next, we tested PDS, a compound developed to be specific for binding GQ DNA (Rodriguez et al. 2008). We pre-incubated 9xTEL with increasing concentrations of PDS and measured CST-PP synthesis in NaCl/KCl or LiCl. PDS did indeed inhibit CST-PP synthesis in Na^+^/K^+^ as we predicted; unexpected, however, was the inhibition in Li^+^ **(Fig. 3A).** The IC50_synthesis_ values were similar: 140 nM and 110 nM, respectively **(Fig. 3B**, **Table 1).** Thus, PDS inhibits synthesis from GQ-forming templates under both GQ-stabilizing and GQ-destabilizing conditions. We next tested whether PDS would inhibit CST-PP synthesis with the 9xnoGG template and were again surprised to find inhibition (**Fig. 3C**), with IC50_synthesis_ similar to that of 9xTEL in both Na^+^/K^+^ and Li^+^ **(Fig. 3D**, **Table 1)** Therefore, PDS inhibits CST-PP synthesis whether the template contains consecutive G’s or not, and in both GQ-stabilizing and GQ-destabilizing conditions.

**FIGURE 3.**
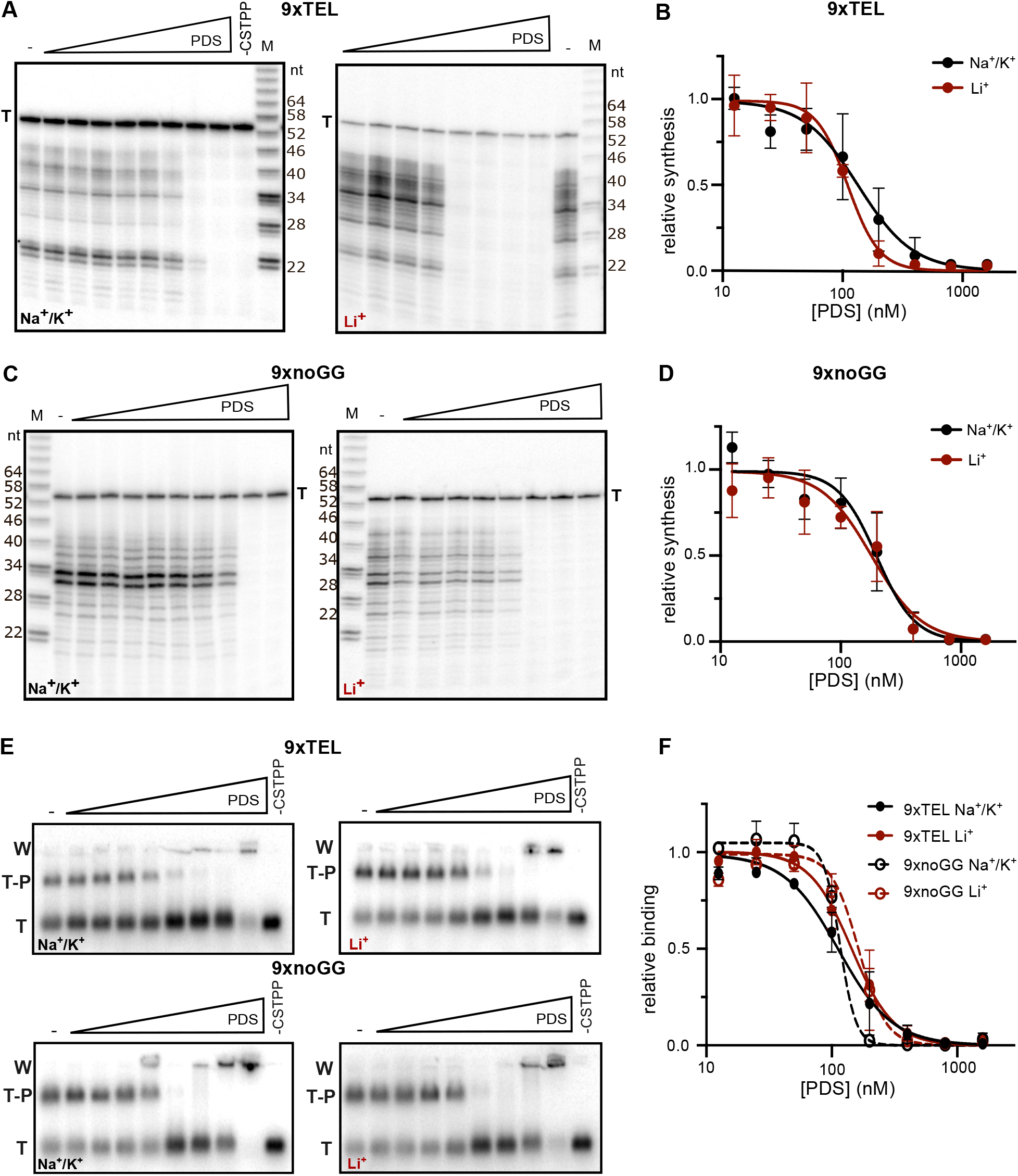
Pyridostatin inhibits C-strand synthesis by inhibiting CST-PP binding to template. (A) Representative gels showing C-strand synthesis in NaCl/KCl (left) or LiCl (right) with trace amount of radiolabeled template (T, 9xTEL) serving as recovery control. Two-fold dilutions of PDS (highest concentration: 1600 nM) were incubated with template prior to addition of CST-PP and NTPs. (B) Plot of relative synthesis intensities with T = 9xTEL at each concentration of PDS in NaCl/KCl or LiCl. Points represent mean values and error bars represent SD for n = 6 (Na^+^/K^+^) or n = 4 (Li^+^) independent experiments. (C) and (D) same as A and B, respectively but T= 9xnoGG. (E) Representative gels of CST-PP binding to radiolabeled template in NaCl/KCl (left) or LiCl (right) following incubation of DNA with increasing concentrations of PDS (two-fold dilutions from 1600 nM. T = Template, T-P = Template-Protein complex, W = Well. (F) Plot of fraction DNA bound at each concentration of PDS normalized to fraction bound with no inhibitor in NaCl/KCl or LiCl. Points represent mean values and error bars represent SD for n = 4 (Na^+^/K^+^) or n = 3 (Li^+^) replicates.

We next evaluated the effect of PDS on CST-PP binding to each of the templates in the two salts **(Fig 3E).** As with synthesis, binding was inhibited in both salts for both templates. The values of IC50_binding_ and IC50_synthesis_ are in agreement **(Fig. 3F**, **Table 1).**, indicating that synthesis inhibition is due to binding inhibition. At high concentrations of PDS, the DNA was aggregated in the well; because this was also seen without protein **(Supplemental Fig. S2C)**, aggregation was not due to CST-PP.

### TMPyP4 and PDS promote secondary structure formation in ssDNA

Considering the lack of inhibition specificity, we wanted to understand how the small molecules were interacting with the templates. Note that although the noGG template sequence was designed to prevent GQ formation, the discovery of the pUG-fold structure in RNA containing 12 or more UG repeats (Roschdi et al. 2022) suggests that one should be cautious in dismissing GQ potential based on sequence alone. To test whether TMPyP4 and PDS were stabilizing GQs or other secondary structures, we performed native 10% polyacrylamide gel electrophoresis of the templates with the small molecules, preparing the DNA-small molecule samples in the two salts as before.

In both the Na^+^/K^+^ and the Li^+^ conditions, 9xTEL mobility subtly increased upon addition of TMPyP4, with a shift too small to be due to GQ stabilization **(Fig. 4A)**. Interestingly, when TMPyP4 was added to 9xnoGG, a second slower migrating band appeared in both salt conditions **(Fig. 4B)**. Although not quite double the apparent molecular weight of the DNA alone, we wondered if the new band represented a dimer. To test this, we mixed 9xnoGG and 5xnoGG with and without drug, reasoning that a heterodimer formed between two different sizes of DNA would have an electrophoretic mobility between those of the two homodimers; thus, appearance of an intermediate band would indicate dimer formation. Upon addition of TMPyP4, a new species was formed in the mixed 9x and 5xnoGG, demonstrating dimer formation **(Supplemental Fig. S3A)**.

**FIGURE 4.**
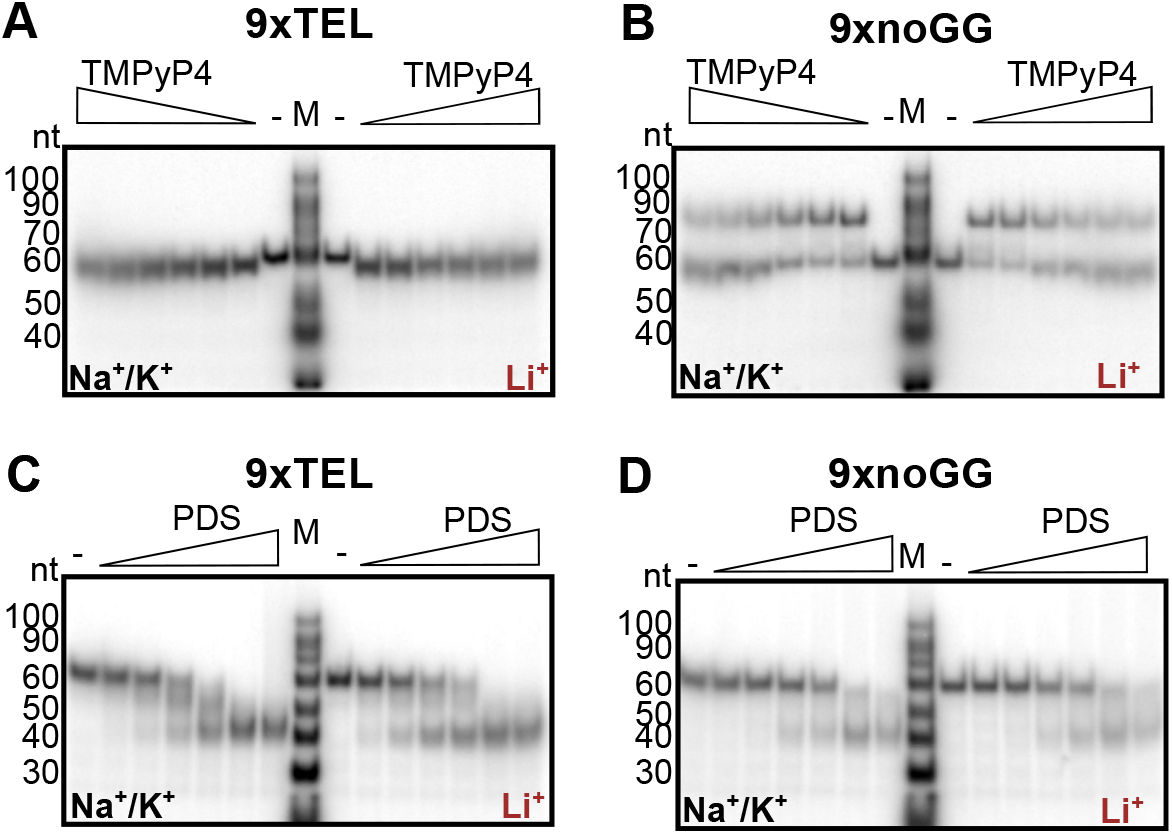
(A) Native gel electrophoresis with 9xTEL DNA incubated with two-fold dilutions of TMPyP4 (highest concentration: 3200 nM) in NaCl/KCl (left half of gel) or LiCl (right half). (B) As in panel A but with 9xnoGG DNA. (C) Native gel of 9xTEL DNA incubated with two-fold dilutions of PDS (highest concentration: 1600 nM) in NaCl/KCl (left half) or LiCl (right half). (D) As in panel C but with 9xnoGG DNA.

We wondered if 9xnoGG might be able to homodimerize without TMPyP4 and rationalized that an intermolecular interaction would be more likely at high DNA concentration. We incubated 25 nM or 5 µM DNA with 5:1 TMPyP4:DNA and performed native gel electrophoresis at 4°C with Na^+^ and K^+^ in the gel and running buffer **(Supplemental Fig. S3B)**.

At 5 µM, 9xnoGG migrated slower, even without TMPyP4; interestingly, in the presence of the small molecule, migration was further slowed, suggesting even higher order oligomerization may occur at high concentrations **(Supplemental Fig. S3B)**. There was no change in 9xTEL migration at high concentration, consistent with intramolecular folding.

PDS had a different influence on the two templates than TMPyP4. The effect on 9xTEL in Na^+^/K^+^ was as expected; a band with higher mobility appeared, indicative of GQ stabilization by PDS. Interestingly the same band appeared in Li^+^ as well (**Fig. 4C**), suggesting that PDS could induce or stabilize a GQ-like structure in 9xTEL, even without GQ-stabilizing cations. To our surprise, PDS likewise increased migration of 9xnoGG, both in Na^+^/K^+^ and in Li^+^ **(Fig. 4D)**, producing a band with the same apparent mobility as the shifted 9xTEL band. Thus, PDS induces or stabilizes a more compact structure in both 9xTEL and 9xnoGG in both Na^+^/K^+^ and Li^+^. Additionally, at 5 µM 9xnoGG, most of the high molecular weight band was shifted into the lower band with the addition of PDS, indicating that the small molecule destabilized the dimeric 9xnoGG relative to the intramolecular structure **(Supplemental Fig. S3B)**.

### TMPyP4 increases stability of ssDNA secondary structures

As the native gels were insufficient to identify the types of secondary structures formed by the ssDNA templates, we turned to circular dichroism (CD) for more information. We first examined 9xTEL in NaCl; with positive peaks at 290 and 240 nm and a negative peak at 260 nm, the spectrum matched published spectra for antiparallel GQ formation **(Fig. 5A)**. We also collected spectra of 9xTEL in Na^+^/K^+^ and in K^+^ only, which were both consistent with hybrid GQ **(Supplemental Fig. S4A-B)**. Then we added TMPyP4 to 9xTEL at a ratio of 10:1 in each salt condition. There was a slight increase in the peak at 260 nm in Na^+^/K^+^, and the K^+^ and Na^+^ spectra did not change **(Supplemental Fig. S4A-B, Fig. 5A).** We performed thermal melts of 9xTEL in Na^+^ with and without TMPyP4, monitoring the signal from the positive peak at 290 nm and the negative peak at 260 nm **(Fig. 5B)**. We found the drug increased the melting temperature (T_m_) by about 10 °C. Thus, 9xTEL DNA in Na^+^ forms an antiparallel GQ that is stabilized by TMPyP4, consistent with previous studies (Martino et al. 2009).

**FIGURE 5.**
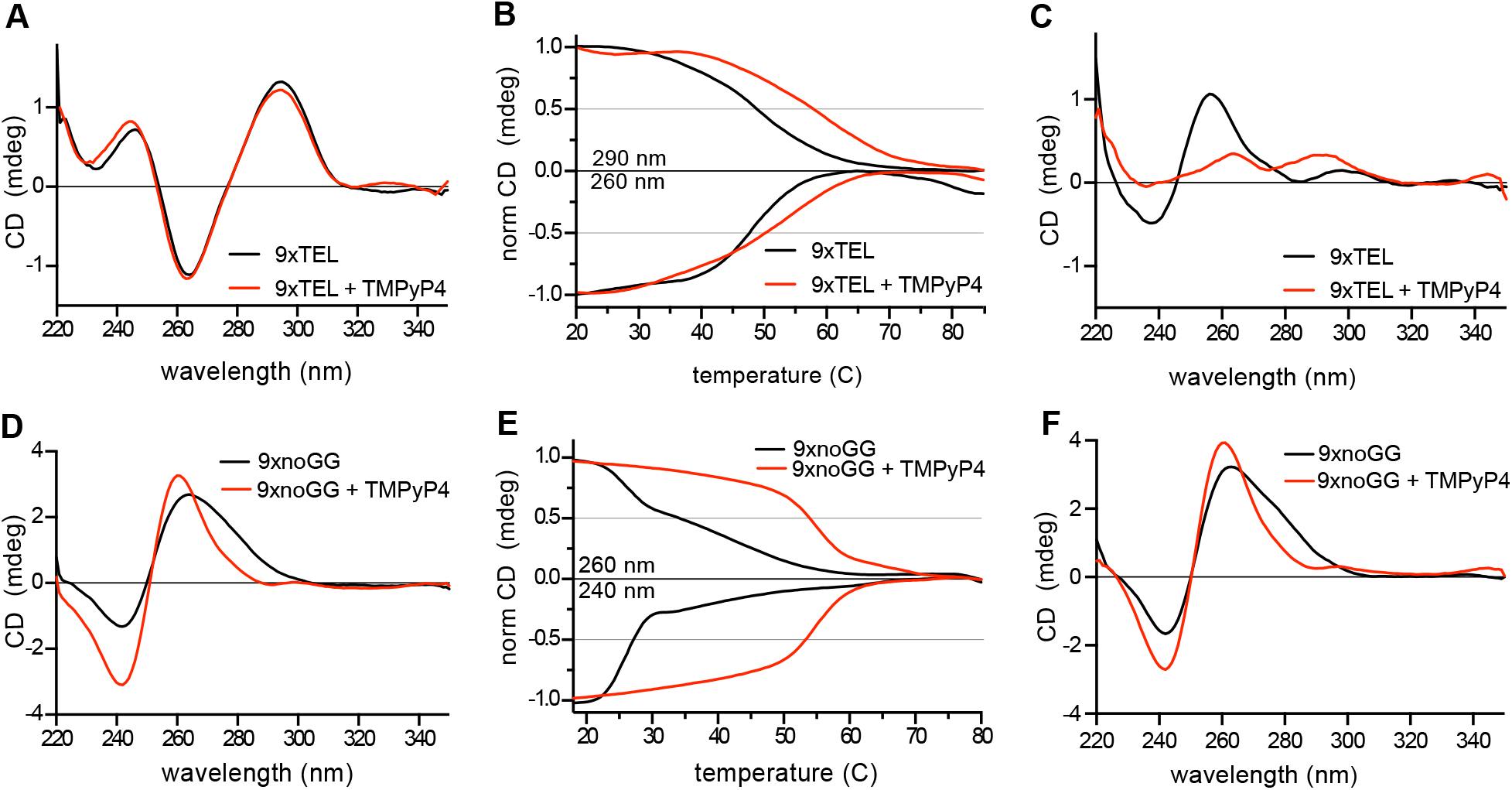
TMPyP4 stabilizes template DNA secondary structure. (A) Circular dichroism spectra of 9xTEL in NaCl with and without 10-fold molar excess TMPyP4. (B). Thermal melting of 9xTEL in NaCl with and without 10-fold molar excess TMPyP4. CD was monitored at 290 nm and 260 nm and normalized between 1 and 0 or between -1 and 0, respectively. (C) CD spectra of 9xTEL in LiCl with and without 10-fold molar excess TMPyP4. (D) Circular dichroism spectra of 9xnoGG in NaCl with and without 10-fold molar excess TMPyP4. (E) Thermal melting of 9xnoGG in NaCl with and without 10-fold molar excess TMPyP4. CD was monitored at 260 nm and 240 nm and normalized between 1 and 0 or between -1 and 0, respectively. (F) Circular dichroism spectra of 9xnoGG in LiCl with and without 10-fold molar excess TMPyP4.

We next acquired spectra for 9xTEL in Li^+^; without small molecule, the spectrum did not resemble parallel, antiparallel, or hybrid GQ spectra **(Fig. 5C)**. We again added 10:1 TMPyP4, and the spectrum gained positive peaks at 290 and 260 nm, indicating formation of a structure that had some CD spectral characteristics of GQ **(Fig. 5C)**. The amplitude of the peaks was reduced relative to those in NaCl **(Fig. 5A)**, likely indicating that much of the DNA remained unstructured at this concentration of inhibitor. The IC50_binding_ and IC50_synthesis_ indicate a higher concentration of TMPyP4 is necessary for inhibition in Li^+^ **(Table 1)**; however, TMPyP4 introduced noise to the spectra, limiting the maximum ratio of small molecule to DNA and preventing spectral analysis at higher concentrations (data not shown). We performed full spectrum thermal melts of 9xTEL in Li^+^ with and without TMPyP4. Since 9xTEL was unstructured in Li^+^ to begin with, there was very little change in the spectrum with increasing temperature (**Supplemental Fig. S4C)**. Due to the melting behavior of 9xTEL with TMPyP4, we were unable to estimate a melting temperature; nonetheless, it was clear that some secondary structure was induced **(Supplemental Fig. S4D)**.

We then collected spectra for 9xnoGG in each salt (Na^+^/K^+^, Na^+^, K^+^, Li^+^) and saw little difference among the conditions **(Fig 5D, Supplemental Fig. S5A)**. There was a broad positive peak at 260 nm with a shoulder around 280 nm, and a negative peak at 240 nm, which could be indicative of A-form double-stranded DNA, parallel G-quadruplex, hairpin DNA, or a mixture of multiple such structures (Kypr et al. 2009; Vorlíčková et al. 2012). Some of the DNA may be dimeric under CD conditions, which require a higher DNA concentration (250 nM DNA used here). Adding TMPyP4 to 9xnoGG in Na^+^ or Li^+^ led to the narrowing of the positive peak at 260 nm, eliminating the shoulder seen in the spectra without drug; the negative peak was not changed **(Fig. 5D)**. The data provided no convincing evidence supporting a GQ structure for this DNA sequence, either in the absence of the presence of TMPyP4.

We performed thermal melts of 9xnoGG in Na^+^ and Li^+^ and monitored the signal at the predominant peaks (positive at 260 nm and negative at 240 nm). The curves were biphasic, with an initial sharp transition at about 26 °C and a minor, broader transition around 40 °C **(Fig. 5E).** We also performed melts with TMPyP4 in both salts and found that it provided substantial stabilization; each had a single transition at 52 °C **(Fig 5E).** Thus, TMPyP4 significantly stabilizes a dimeric form of 9xnoGG.

### Pyridostatin stabilizes GQ structure of single-stranded telomeric DNA

We next gathered spectra with 10:1 PDS:DNA, starting with 9xTEL. As has been described for shorter oligonucleotides (Marchand et al. 2015), PDS induced a shift from hybrid to antiparallel GQ in Na^+^/K^+^ **(Fig. S6A)**. There was no change in the spectra with and without PDS in Na^+^, where 9xTEL is already in the antiparallel conformation **(Fig. 6A).** In Li^+^, where 9xTEL is unstructured without drug, addition of PDS induced a structural shift indicative of antiparallel GQ **(Fig. 6B).** The spectra of 9xTEL with PDS in each salt were remarkably indistinguishable **(Supplemental Fig. S6B),** indicating that PDS permits or facilitates antiparallel GQ-formation of 9xTEL in either stabilizing or destabilizing cations.

**FIGURE 6.**
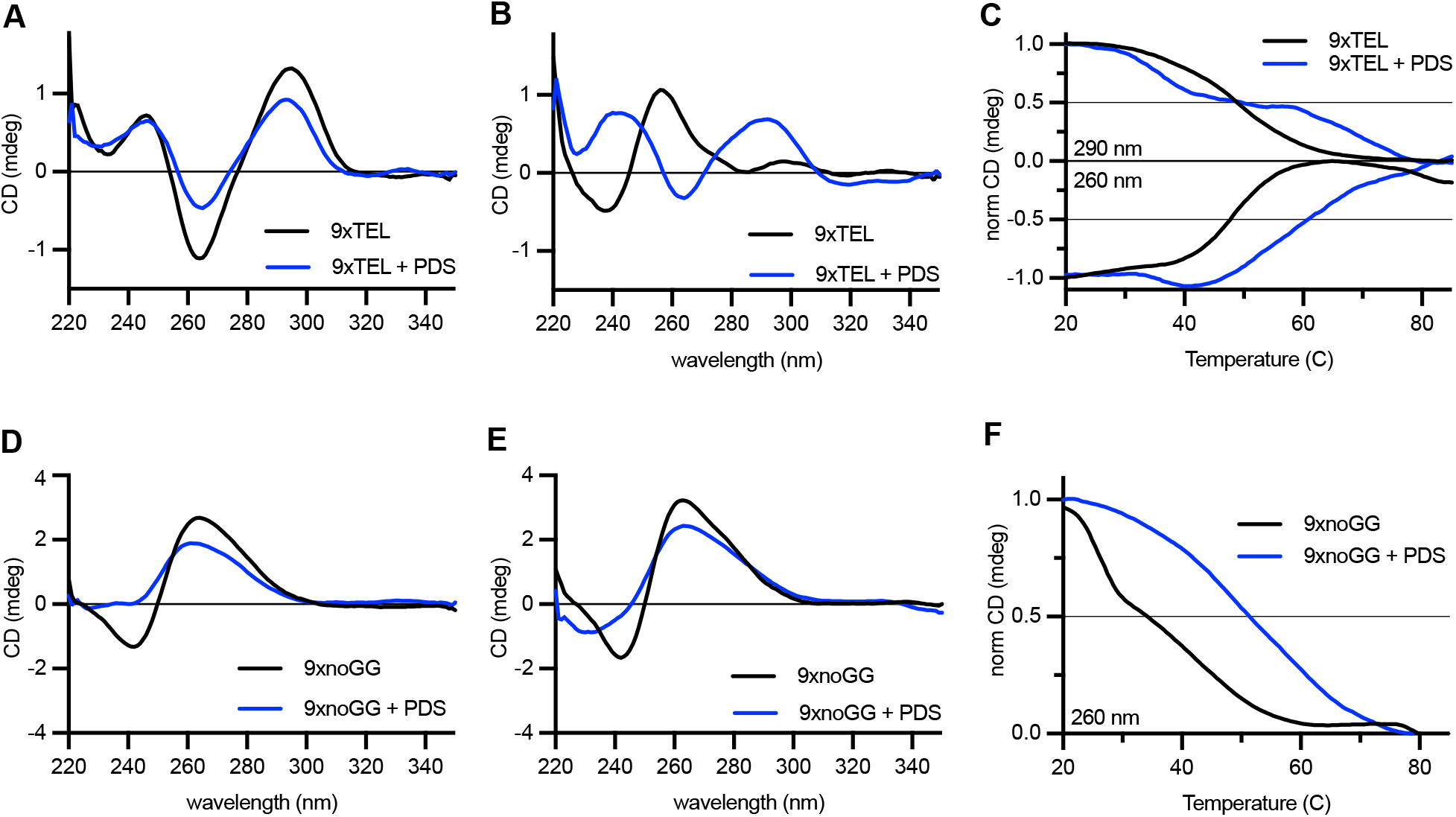
Pyridostatin stabilizes template DNA secondary structure. (A) Circular dichroism spectra of 9xTEL in NaCl with and without 10-fold molar excess PDS. (B) CD spectra of 9xTEL in LiCl with and without 10-fold molar excess PDS. (C) Thermal melting of 9xTEL in NaCl with and without 10-fold molar excess PDS. CD was monitored at 290 nm and 260 nm and normalized between 1 and 0 or between -1 and 0, respectively. (D) Circular dichroism spectra of 9xnoGG in NaCl with and without 10-fold molar excess PDS. (E) CD spectra of 9xnoGG in LiCl with and without 10-fold molar excess PDS. (F) Thermal melting of 9xnoGG in NaCl with and without 10-fold molar excess PDS. CD was monitored at 260 nm and normalized between 1 and 0.

We then melted 9xTEL with PDS. While 9xTEL alone had a monophasic melt, with PDS there were two transitions at 290 nm **(Fig. 6C)**. Additionally, distinct peaks persisted even above 90°C, unlike in 9xTEL alone **(Supplemental Fig. S6C)**. We deduced that the complex could not be entirely melted; therefore, although it was not appropriate to calculate a melting temperature, we concluded that there was substantial stabilization of GQ secondary structure by PDS. We also performed full spectrum thermal melts of 9xTEL with PDS in Li^+^, which likewise indicated considerable stabilization **(Supplemental Fig. S6D)**.

Unlike 9xTEL, PDS did not shift 9xnoGG into an antiparallel GQ, but it did induce a conformation change **(Fig 6D, E**). In both Na+ and Li+ conditions, the two peaks were attenuated with respect to the no-drug spectrum, and the negative peak at 240 nm was flattened. Melts of 9xnoGG with PDS in both salts were monophasic when monitoring the positive peak at 260 nm, but there was no significant change in the signal at 240 nm **(Fig 6F, Supplemental Fig. S6E)**. The T_m_ was slightly higher in Na^+^ than in Li^+^. Nonetheless, the data indicate that PDS stabilizes some secondary structure in 9xnoGG in all cations tested.

## DISCUSSION

Here we show that GQ-stabilizing molecules TMPyP4 and PDS inhibit telomeric C-strand synthesis by CST-PP by interacting with the single-stranded DNA templates and thereby preventing binding of the enzyme. Both molecules also inhibit telomerase *in vitro*, but under our standard reaction conditions their effect on CST-PP was stronger (lower IC50) than on telomerase. Thus, effects of these compounds on telomere biology in living cells may very well be due to perturbation of G-strand synthesis, C-strand synthesis, and very likely other telomeric processes. Due to the additional genomic roles CST-PP plays, such as 3’ fill-in prior to double-strand break repair, inhibition of CST-PP is likely to have implications beyond the telomere.

This study also demonstrated unexpected interactions between these small molecules and single-stranded DNA in non-GQ-stabilizing conditions. TMPyP4 inhibited CST-PP better in the Na^+^/K^+^ condition than in Li^+^, but with only a 4-fold difference in IC50. PDS inhibited synthesis and binding equally in both conditions, and CD spectroscopy demonstrated that although 9xTEL is unfolded in lithium ion, addition of PDS causes GQ formation. This suggests that PDS not only binds to pre-existing GQs but also drives the structure in otherwise unfavorable conditions. Similarly, another GQ-stabilizing molecule, telomestatin, has been shown to induce GQ-formation without monovalent cations (Kim et al. 2003; Gao et al. 2017). Mass spectrometry has also revealed that PDS binding induces cation ejection and quartet rearrangement (Marchand et al. 2015; Lecours et al. 2017). Taken together, these results indicate that traditional methods of GQ destabilization are not sufficient to inhibit GQ formation in PDS.

We also showed that the small molecules stabilized secondary structure in a DNA sequence without consecutive G’s, designed to prevent GQ formation. TMPyP4 stabilized a dimeric form of 9xnoGG, increasing the melting temperature by over 20 °C. Others have reported TMPyP4-induced dimerization of shorter TEL-containing sequences under certain conditions (Gao et al. 2017), although we did not see dimerization of 9xTEL in this study. The lack of specificity of TMPyP4 for GQ DNA could have been anticipated, as others have found that this compound binds many nucleic acids, including double-stranded DNA (Ren and Chaires 1999; De Cian et al. 2005). PDS also interacted with 9xnoGG, and while it is not clear what sort of structure is formed, the increased melting temperature over unliganded 9xnoGG indicates that it is quite stable. The IC50_inhibition_ and IC50_binding_ of PDS for 9xTEL and 9xnoGG are also the same, suggesting that PDS has little specificity for the canonical GQ structure over the unknown structure.

A sequence with no consecutive G’s seems unlikely to form a G-quadruplex; however, recent identification of new noncanonical GQ structures, including an RNA poly-UG (pUG) fold (Roschdi et al. 2022) and DNA GQs containing large loops, bulges, or guanine vacancies (for reviews on noncanonical GQs, see refs (Banco and Ferré-D’Amaré 2021; Jana et al. 2021), demonstrates that understanding the requirements for GQ formation is still developing. Although computational methods can identify potential GQ-containing sequences within the genome, they face several challenges. First, in cells these sequences may be effectively prevented from forming GQs by the complementary strand of the DNA double helix, except at transiently single-stranded regions such as open promoters and replication forks, and at telomeres. In the other direction, computational predictions may miss intermolecular GQs or noncanonical GQs, including those that contain gaps, bulges, and loops. Thus, GQ-stabilizing small molecules have been employed as a biochemical method of identification or validation; pyridostatin-based GQ identification found over 700,000 GQ-forming sequences in human cells, approximately 70% of which were not predicted by sequence alone (Chambers et al. 2015). It is unclear what proportion of these sites would exist as a GQ without small molecules, which may induce their formation. Similarly, if in fact the 9xnoGG DNA forms some sort of GQ, it appears to be either an unstable GQ that is stabilized upon addition of PDS or a structure that is induced by PDS. Such possibilities should be examined upon PDS-mediated identification of unpredicted GQs prior to assigning biological significance.

We did not investigate the effect the DNA-binding compounds might have on other telomeric or nontelomeric protein components. However, considering that CST-PP was effectively prevented from binding to DNA by TMPyP4 and PDS under all conditions tested and considering the number of potential interaction sites in the genome, it is likely that many other DNA-binding proteins would also be affected. For example, replication protein A (RPA) shares both structural and functional similarities with CST, including unfolding GQs (Barbour and Wuttke 2023), and at least one study has reported RPA inhibition due to TMPyP4 stabilization of DNA *in vitro* (Prakash et al. 2011). Pyridostatin has also been shown to inhibit binding of POT1 *in vitro* (Rodriguez et al. 2008) and transcription factors in cells (Spiegel et al. 2021). Although the helicase Pif1 reportedly resolved GQs stabilized by the small molecule BRACO19 *in vitro* (Zhou et al. 2014) and overexpression of Pif1 partially rescued PDS-induced phenotypes in cultured primary neurons (Moruno-Manchon et al. 2020), other studies have described helicase inhibition by GQ stabilizers *in vitro* (Wu and Maizels 2001; Mendoza et al. 2015; Maleki et al. 2019; Wu et al. 2019) and in cells (Wu et al. 2019). Thus, it’s still unclear whether small molecule stabilized GQs are effectively resolved in cells. The present study also raises the question of reversibility; even after reaching boiling, PDS-bound 9xTEL was not fully denatured.

With questions of specificity, unexpected sequences/structures affected, questionable reversibility and resolvability, and an ever-growing list of proteins and processes affected by TMPyP4 and PDS, extra caution should be used when employing these small molecules, particularly in cells. Furthermore, these compounds bind to RNA, so RNA-protein interactions are likely to be affected. It certainly remains possible that other GQ-stabilizing molecules exhibit higher specificity for GQ structures than TMPyP4 and PDS. The sort of analysis of inhibitor specificity performed here and previously by others should be useful in assessing additional small molecule inhibitors.

## MATERIALS AND METHODS

### CST-Polymerase α-primase purification

CST-Polymerase α-primase was purified from HEK 293T/17 cells (CRL-11268, ATCC) as previously described(Zaug et al. 2021, 2022). Briefly, cells were transfected with 3xFLAG-CTC1, Myc-STN1, and HA-TEN1 at a ratio of 1:1:1, expanded and grown for 48 hours, and collected. The complex was purified by successive anti-FLAG and anti-HA immunoprecipitation. Endogenous Polα-primase copurifies with CST (Zaug et al. 2022). The purity of the complex was verified by silver staining and concentrations were determined by Western blot analysis (Zaug et al. 2022).

### Small molecule inhibitors

TMPyP4 (sc-204346, Santa Cruz Biotechnology) and pyridostatin (SML2690, Sigma-Aldrich) were resuspended in 100% DMSO at 10 mg/mL and stored at -20 °C.

### Oligonucleotides

DNA oligonucleotides were purchased from Integrated DNA Technologies (IDT) and resuspended in water. 5’-end radiolabeled DNA was generated with T4 polynucleotide kinase (M0201L, NEB) and [γ-^32^P]ATP (NEG035C005MC, PerkinElmer) according to manufacturer’s standard protocol.

### Telomerase inhibition

Telomerase reactions (20 uL) contained 2 nM telomerase and 50 nM 3xTEL DNA in 1x telomerase activity buffer (50 mM Tris-HCl pH 8.0, 2 mM MgCl_2_, 1 mM spermidine, and 5 mM β-mercaptoethanol, 75 mM KCl, and 50 mM NaCl), and dNTPs (0.5 mM dGTP, 0.5 mM dTTP, 0.33 μM [α-^32^P]dATP (300 Ci mmol^-1^), 3.3 μM unlabelled dATP), the indicated concentrations of TMPyP4 or pyridostatin, and 10% DMSO. All components were mixed and the samples were incubated at 30 °C for 1 h. Reactions were quenched with 100 µL LC stop mix (3.6 M ammonium acetate, 0.2 mg ml^−1^ glycogen, and a 16-nt ^32^P-labelled loading control) and ethanol precipitated. Pellets were dried and dissolved in equal volumes water and loading dye (95% formamide, 20 mM EDTA, and 0.05% each bromophenol blue and xylene cyanol), then boiled and run on a sequencing-style 10% acrylamide, 7 M urea, 1× TBE gel. The bromophenol blue dye was run to the bottom of the gel (about 1.75 h). The gel was removed from the glass plate using Whatman 3-mm paper and then dried at 80 °C under vacuum for 15–30 min. Gels were then exposed to a phosphorimager screen, which were imaged on a Typhoon FLA9500 scanner (GE Lifesciences) and analyzed by ImageQuant TL v.8.1.0.0 (GE Lifesciences). The intensity of synthesis was normalized to the intensity of the loading control in the same lane. Relative activity was calculated by dividing normalized synthesis at each inhibitor concentration by the normalized synthesis without inhibitor. The results were plotted with each point representing n = 3 independent replicates; a four-parameter logistic curve with bottom constrained to 0 was fitted to determine IC50s (Prism).

### CST-PP synthesis inhibition

Reactions (10 μL) contained 25 nM CST-PP and 25 nM single-stranded template DNA in 1x reaction buffer (50 mM Tris-HCl pH 8.0, 2 mM MgCl_2_, 1 mM spermidine, and 5 mM β-mercaptoethanol), 1x Na^+^/K^+^ (75 mM KCl and 50 mM NaCl) or 1x Li^+^ (125 mM LiCl) salts, rNTPs (0.2 mM each CTP, UTP, ATP), and dNTPs (0.5 mM dCTP, 0.5 mM dTTP, 0.33 μM [α-^32^P]dATP (300 Ci mmol^-1^), 0.29 μM unlabelled dATP), the indicated concentrations of TMPyP4 or PDS, and 0.2% DMSO.

DNA templates and small molecule dilutions were prepared fresh immediately prior to each use. A small proportion of the template DNA was 5’ radiolabeled (typically about 0.1 – 0.5 nM, 5000 counts per minute per reaction) to serve as a recovery and loading control and internal size marker. CST-PP generated more product with 9xnoGG in Na^+^/K^+^ than in other conditions, so in that condition 5x more radioactive template was used (keeping total DNA at 25 nM) so that the control signal was not overwhelmed by product signal. DNA templates in buffer and designated salt were placed in 95 °C heat block for 5 min and cooled at room temperature for 30 min. Dilutions of PDS or TMPyP4 in 1% DMSO (or 1% DMSO only for no inhibitor condition) were added (final 0.2% DMSO), and mixtures were incubated in 30 °C water bath for 30 min. CST-PP was added and mixtures were returned to water bath for 30 min. NTPs were added and the reaction proceeded at 30 °C for 1 hour; 50 µL stop mix (3.6 M ammonium acetate and 0.2 mg ml^−1^ glycogen) was then added.

Quenched reactions were cleaned up using the Oligo Clean & Concentrator kit (D4060, Zymo) and eluted in water. An equivalent volume of 2x loading dye (95% formamide, 20 mM EDTA, and 0.05% each bromophenol blue and xylene cyanol) was added. Samples were boiled for 5 min, then loaded on a 10% acrylamide, 7 M urea, 1x TBE gel. Molecular weight ladders were generated by telomerase as above (no small molecule). Gels were run at 30 W until the bromophenol blue was about 2 inches from the bottom (∼1-1.5 hours) and dried on Whatman 3-mm paper at 80 °C under vacuum for 15-30 min. Gels were then exposed to a phosphorimager screen, which was imaged with Amersham Typhoon scanner 5 and analyzed using ImageQuant. The intensity of synthesis was normalized to the intensity of the loading control in the same lane. Relative synthesis was calculated by dividing normalized synthesis at each inhibitor concentration by the normalized synthesis without inhibitor. The results were plotted with each point representing the mean of n ≥ 3 independent replicates; a four-parameter logistic curve with bottom constrained to 0 was fitted to determine IC50s (Prism).

### Binding inhibition

Binding reactions were prepared identically to synthesis reactions except glycerol (10% final) was added in place of the NTP mix. DNA and small molecule were incubated for 30 min at 30 °C, then CST was added and incubation continued for another hour. Samples were loaded onto a 1x TBE 0.7% agarose gel and run at 70 V for 75 min at 4 °C. The gels were placed on Hybond N+ atop Whatman 3-mm paper and dried under vacuum at 80 °C for 1 hour. They were then exposed to a phosphorimager screen, which was imaged using an Amersham Typhoon scanner 5. Binding was analyzed using ImageQuant; fraction bound was determined by dividing the number of counts in the bound band by the total number of counts in the lane, not including the well. Relative binding was calculated by dividing fraction bound at each inhibitor concentration by the fraction bound without inhibitor. Results were plotted and analyzed as above.

### DNA native gels

DNAs for native gels were prepared as above with glycerol but without CST or NTPs. Following incubation at 30 °C for 30 min, the samples were loaded on 10% polyacrylamide gels containing 1x TBE or containing 0.5x TBE and 0.5x Na^+^/K^+^ as indicated. The ladder was a 1:1 mix of 5’ radiolabeled 10/60 ladder (51-05-15-01, IDT) and 20/100 ladder (51-05-15-02, IDT) in 1x buffer, 1x salt, 10% glycerol, and 0.025% bromophenol blue. Gels were run at 7 W submerged in 1xTBE (or 1/2x TBE and 1/2x Na^+^/K^+^ where indicated) until the bromophenol was just more than halfway through the gel. Gels with radiolabeled samples were dried on Whatman 3-mm paper and under vacuum at 80 °C for 1 hour and exposed to a phosphorimager screen which was imaged with Amersham Typhoon scanner 5.

### Circular dichroism

CD samples contained 250 nM DNA in 75 mM NaCl and 50 mM KCl (Na^+^/K^+^), or 125 mM NaCl, KCl, or LiCl, 0.2x reaction buffer (10 mM Tris-HCl pH 8.0, 0.4 mM MgCl_2_, 0.2 mM spermidine, and 1 mM β-mercaptoethanol), and 0.02% DMSO; those with inhibitors also included 2.5 µM TMPyP4 or PDS. Baseline control samples consisted of 1x corresponding salt and 0.2x reaction buffer.

Samples containing DNA, salt, and buffer were placed in 95 °C heat block for 10 min and cooled at room temperature for 30 min. PDS, TMPyP4, or DMSO was added and samples were incubated at 30 °C for 30 min. 1.8 mL was added to a 10 mm pathlength cuvette and loaded into a Chirascan Plus Circular Dichroism and Fluorescence Spectrometer (Applied Photophysics). Spectra were collected between 360 nm and 220 nm with a step size of 1 nm. Three repeats were averaged and smoothed with a second order polynomial with 10 neighbors on each side (Prism).

Continuous ramped thermal melts were performed at the indicated wavelengths with a 2 nm bandwidth. The temperature was increased from 20 °C to 90 °C at a rate of 1 °C/min with a time-per-point of 12 sec. Positive peaks were normalized to 1 at the lowest temperature and negative peaks were normalized to -1 at the lowest temperature. Spectra were smoothed as above.

Stepwise thermal melts were performed where continuous ramps were unsuitable due to anomalous melting behavior. Full spectra were collected between 360 nm and 220 nm with a step size of 1 nm every 2 °C with a settling time of 180 sec. For clarity, every other spectrum was omitted from figures (i.e. a spectrum is shown for every 4 °C increase in temperature). Spectra were smoothed as above.

## SUPPLEMENTAL MATERIAL

Supplemental material is available for this article.

## ACKNOWLEDGMENTS

We thank Art Zaug for advice on CST-PP purification and for samples of telomerase and CST-PP and Shankar Balasubramanian for discussions about GQ formation assays. Special thanks to Annette Erbse for advice on CD spectrometry and interpretation of spectra. We thank the Shared Instrument Pool (RRID: SCR_018986) of the Department of Biochemistry, CU-Boulder for the use of the CD spectrometer. The CD is funded by NIH Shared Instrumentation grant S10RR028036. T.R.C. is an investigator of the Howard Hughes Medical Institute.

## Author contributions

K.J. and T.R.C. designed the experiments. J.S. performed the initial studies of telomerase and CST-PP inhibition, which were replicated and extended by K.J. K.J. designed and performed the experiments assessing small molecule-DNA interactions by gel electrophoresis and CD. K.J. and T.R.C. wrote the manuscript with input from all authors.

